# Rationale design of unrestricted pRN1 derivatives and their application in construction of a dual plasmid vector system for *Saccharolobus islandicus*

**DOI:** 10.1101/2023.10.26.564113

**Authors:** Pengpeng Zhao, Xiaonan Bi, Xiaoning Wang, Xu Feng, Yulong Shen, Guanhua Yuan, Qunxin She

**Affiliations:** CRISPR and Archaea Biology Research Center, State Key Laboratory of Microbial Technology and Microbial Technology Institute, Shandong University, Qingdao 266237, China

**Keywords:** CRISPR-Cas, Cryptic plasmids, CRISPR escape mutations, *Saccharolobus-E. coli* shuttle vectors, dual plasmid vectors, archaeal genetics

## Abstract

*Saccharolobus islandicus* REY15A represents one of the very few archaeal models with versatile genetic tools, including efficient genome editing, gene silencing and robust protein expression systems. However, plasmid vectors constructed for this crenarchaeon thus far are solely based on the pRN2 cryptic plasmid. Although this plasmid co-exists with pRN1 in their original host, early attempts to test pRN1-based vectors consistently failed to yield any stable host-vector system for *Sa. islandicus*. Herein we identified a putative target sequence in *orf904* encoding a putative replicase on pRN1 (TargetN1). Mutated targets were then designed (N1a, N1b, N1c) and tested for their capability of escaping from the host CRISPR immunity by using plasmid interference assay. This revealed that the original target triggers the CRISPR immunity in this archaeon whereas all three mutated targets do not, indicating that all designed target mutations evade the host immunity. These mutated targets were then incorporated into *orf904* individually, yielding corresponding mutated pRN1 backbones with which shuttle plasmids were constructed (pN1aSD, pN1bSD and pN1cSD). *Sa. islandicus* transformation revealed that pN1aSD and pN1bSD were functional shuttle vectors, but pN1cSD lost the capability of replication. In addition, pRN1-based and pRN2-based vectors were stably maintained in the archaeal cells either alone or in combination, and this yielded a dual plasmid system for genetic study with this important archaeal model.

**Impact statement:** When pRN1 was employed for vector construction in *Saccharolobus islandicus* REY15A, pRN1-derived vectors are not stable in this archaeon. Here we show that pRN1 orf904 encoding a putative replicase on pRN1 carries a DNA segment to be targeted by the host I-A CRISPR system. By designing mutated target sequences that evade the CRISPR immunity, efficient plasmid vectors were obtained with mutated pRN1 backbones. This strategy could be applied in developing host-vector systems for other microorganisms with plasmids or viruses carrying CRISPR target sequences. Moreover, the resulting dual vector system would facilitate genetic studies with this crenarchaeal model.

## INTRODUCTION

Archaea were first discovered as the third domain of life by Carl Woese and coworkers in 1970s in their pioneer phylogenetic study of microorganisms based on the small-subunit ribosomal RNA sequences (1, 2). This classification was then confirmed by subsequent genomics studies of several archaea and bacteria since genome sequence analysis revealed that Archaea share information-processing machineries with Eukarya, such as proteins/enzymes responsible for DNA replication and RNA transcription (3). On the other hand, Archaea also exhibit unique cell biology (4). Over 20000 archaea are known, falling into 30 phyla (5). In particular, the discovery of archaea belonging to the Asgard superphylum represents one of most exciting research breakthroughs in the contemporary biology: these archaea are probably ancestors to eukaryotes since they share hundreds of proteins involved in cellular processes such as membrane trafficking with eukaryotes (6, 7). Therefore, archaeal models are not only important for investigation of novel biological principles in their own right but also for the origin of eukaryotes and the evolution of eukaryotic biological features (4, 8, 9).

Currently, archaeal genetic and functional studies are still limited to a few model organisms belonging to the two classical phyla, i.e., *Euryarchaeota* and *Crenarchaeota* (10), among which *Sulfolobus acidocaldarius* and *Saccharolobus islandicus* of the latter phylum represent the genetically tractable archaeal models (11, 12, 13) that are more closely related to Asgard archaea, the predicted most recent ancestor of eukaryotes. Since their discovery by metagenomic analysis of marine samples in 2015 (14), great cultivation efforts only yielded two Asgard cultures and both organisms grow extremely slowly (15, 16), a feature that precludes them to be developed as a genetic model in a near future. As a consequence, other models have to be employed for genetic and functional characterization of the eukaryotic signature proteins identified in Asgard archaea. In this regard, *Sulfolobales* organisms, such as *S. acidocaldarius* and *Sa. islandicus*, serve as important models for these investigations since they are more closely related to Asgard archaea than *Euryarchaeota*.

Versatile genetic tools are available for *S. acidocaldarius* and *Sa. islandicus* (11, 12, 13, 17). However, vectors constructed for the former are based on pRN1 (18) whereas those constructed for the latter are based on pRN2 (19). Interestingly, both cryptic plasmids co-exist in *Sa. islandicus* REN1H1, the original host (20), suggesting that they could provide compatible archaeal origins of replication for developing dual vector system in *Sa. islandicus*.

*Sa. islandicus* REY15A was isolated from an enrichment culture generated from hotspring samples collected in the Reykjavík region of Iceland (21) and it grows optimally at 78 °C, pH 3.5 (22). This archaeon has been a model for investigation of antiviral mechanisms by different CRISPR-Cas systems, including I-A and III-B subtypes that employ ribonucleoprotein complexes of multiple subunits to mediate nucleic acid interference in a small RNA-guided fashion (23, 24, 25). In archaea biology research, studies with this crenarchaeon have contributed to the understanding of novel DNA damage-responsive regulation networks (26, 27, 28), the archaeal ESCRT cell division system (29, 30) and cell cycle regulation (31, 32). At present, versatile genetic tools have already been developed for this model, including conventional and novel schemes of genetic manipulations (19, 33), highly efficient expression systems (34, 35) and CRISPR-based mutagenesis and gene silencing (36, 37, 38). Herein, we aimed to test a dual host-vector system for *Sa. islandicus* REY15A to further enrich its genetic toolbox. CRISPR escape mutations were designed on pRN1 and tested for their capability of overcoming the CRISPR restriction to the plasmid, yielding highly efficient two plasmid vector systems for this model crenarchaeon.

## RESULTS AND DISCUSSION

### Spacer L2S56 in the CRISPR array of *Sa. islandicus* REY15A facilitates the I-A immunity but fails to trigger the immune responses from III-B systems

In our first attempts to construct shuttle vectors for *Sa. islandicus* REY15A, both pRN1 and pRN2 plasmids, the cryptic plasmids that coexist in the *Sa. islandicus* REN1H1 (21, 39, 40), were individually employed as an archaeal origin of replication in combination with the *E. coli* cloning vector pGEM3z, and resulting plasmid constructs were then tested by transformation of *Sa. islandicus* E233S, the genetic host derived from REY15A (41). It was reported that, while pRN2-derived plasmids scored a high transformation rate and yielded true transformants (19), pRN1-based vectors yielded only very few colonies from which plasmids were apparently absent (41). Since CRISPR arrays often carry spacers matching various plasmids in *Sulfolobales* (42, 43), we suspected that the genome of *Sa. islandicus* REY15A might carry spacer that matches a sequence in pRN1 but not on pRN2. After the determination of the complete genome sequence of *Sa. islandicus* REY15A (44), it became clear that the host genome indeed carries a spacer homologous to a DNA segment in pRN1 plasmid. It is the 56^th^ spacer in Locus 2 CRISPR array (L2S56), and the matching DNA segment is within the coding sequence of orf904 and it is 44 bp long, showing two mismatches that are individually positioned at 20 nt and 35 nt of the target (TargetN1, Fig. 1A). Since the archaeal host carries one I-A and two III-B CRISPR-Cas systems (Fig. S1), we analyzed if TargetN1 could be recognized for destruction by these host CRISPR immune systems. Previous works showed that a functional protospacer adjacent motif (PAM) at the 5’-flanking position is required for eliciting I-A immunity in *Saccharolobus* (45, 46, 47). Indeed, TargetN1 is immediately downstream of the 5’-CCG-3’ trinucleotide (Fig. 1A), which is one of the strong PAMs for I-A systems. Thus, this DNA sequence is very likely to be targeted for destruction by the host I-A CRISPR-Cas. Furthermore, TargetN1 is on the template strand of *orf904*, meaning that mRNAs of the replicase gene are complementary to the crRNAs of L2S56. Since there are four mismatches between the upstream protospacer-flanking sequence (PFS, 5’-ATCCG-3’) and the 5’-GAAAG-3’ repeat tag of crRNAs in *Sa. islandicus* REY15A (48), upon transcription, mRNAs of *orf904* contain a cognate target RNA sequence that could trigger immune responses of the host III-B Cmr systems (Fig. S1), both of which are active in antiviral defense (49, 50, 51, 52). On the other hand, L2S56 exhibits 8 mismatches to the corresponding segment in pRN2, including a non-PAM motif (5’-CAT-3’, Fig. 1A), and this would render the host CRISPR immunity ineffective against pRN2 in *Sa. islandicus* REY15A.

**Figure 1.**
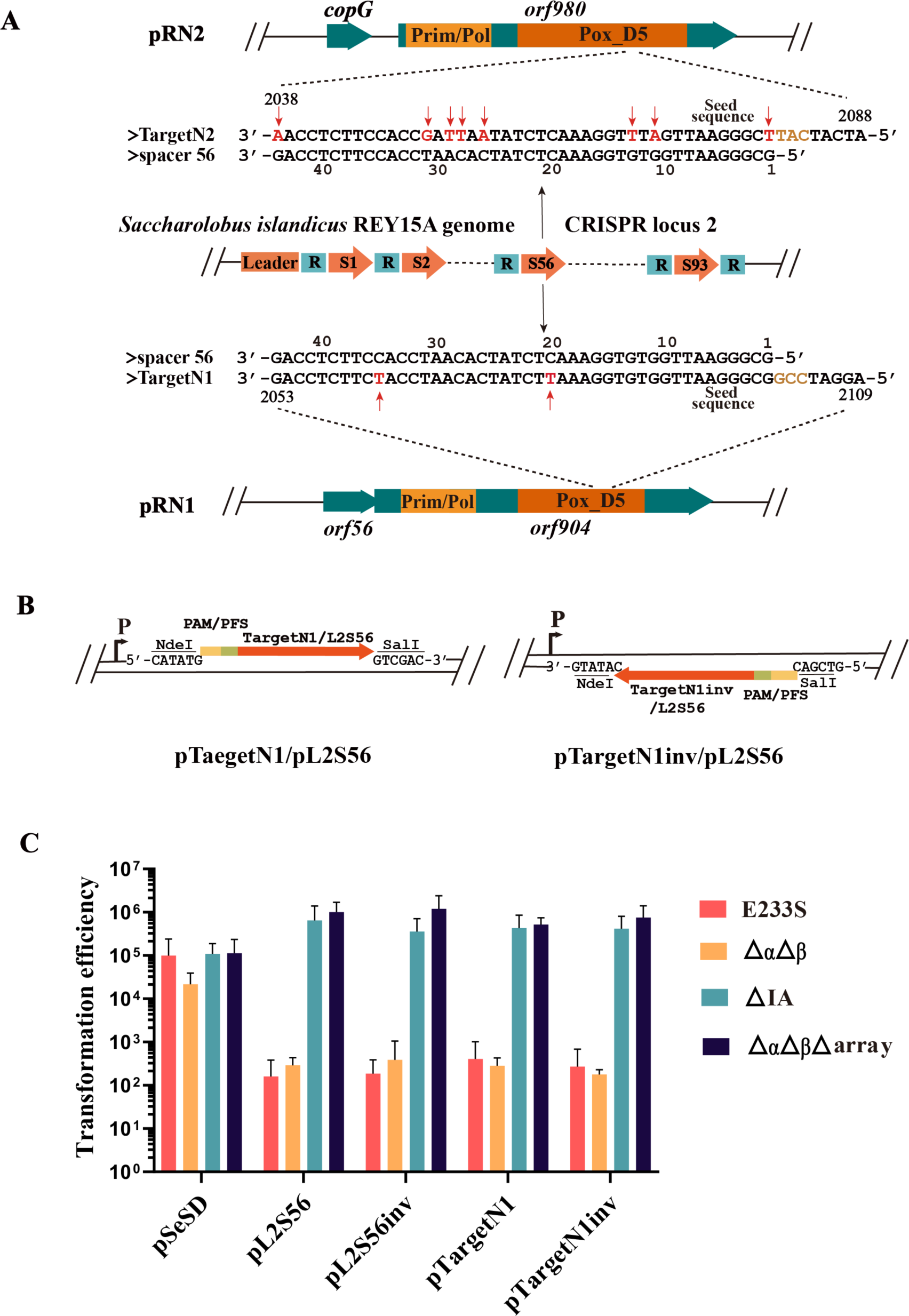
Restriction of pRN1-derived plasmids by CRISPR-Cas systems in *S. islandicus* REY15A. (A) Sequence alignment of TargetN1/TargetN2 and L2S56. L2S56: spacer 56 in CRSPR locus 2 of the host genome; TargetN1: the pRN1 sequence matching the L2S56 spacer. TargetN2: the pRN2 sequence showing similarity to the L2S56 spacer. Mismatches are highlighted in red and indicated with red arrows. The trinucleotides at the PAM position are highlighted in brown. (B) Schematic of interference plasmids. For the artificial L2S56 protospacer (L2S56) and TargetN1, they both contain the 5’ flanking sequence 5’-AGGATCCG-3’ of the pRN1 target. (C) Testing of CRISPR immunity using the two sets of interference plasmids in REY15A. Four strains were employed for transformation, including E233S: the wild-type host; ΔIA: derived from E233S, lacking the *cas* gene module of IA CRISPR-Cas; ΔαΔβ: derived from E233S lacking Cmr-α and Cmr-β modules of *cas* genes; ΔαΔβΔarray: derived from ΔαΔβ with both CRISPR arrays also deleted.

To test the above assumptions, target sequences were designed for L2S56 (ProtoL2S56) and TargetN1, including the 5’-CCG-3’ PAM and 44 bp target sequence (Fig. 1B). The resulting target modules were cloned to pSeSD in both orientations, yielding two types of plasmids, one with the same orientation as the promoter, i.e., pTargetN1 and pL2S56, whereas the other in the reverse orientation, namely pTargetN1inv and pL2S56inv (Fig. 2A). Theoretically, both types of target plasmid are to be recognized by the I-A system, but only the plasmids carrying the inverted target sequence can produce cognate target RNAs that activate the immune responses of III-B CRISPR systems in this archaeon (53, 54, 55).

**Figure 2.**
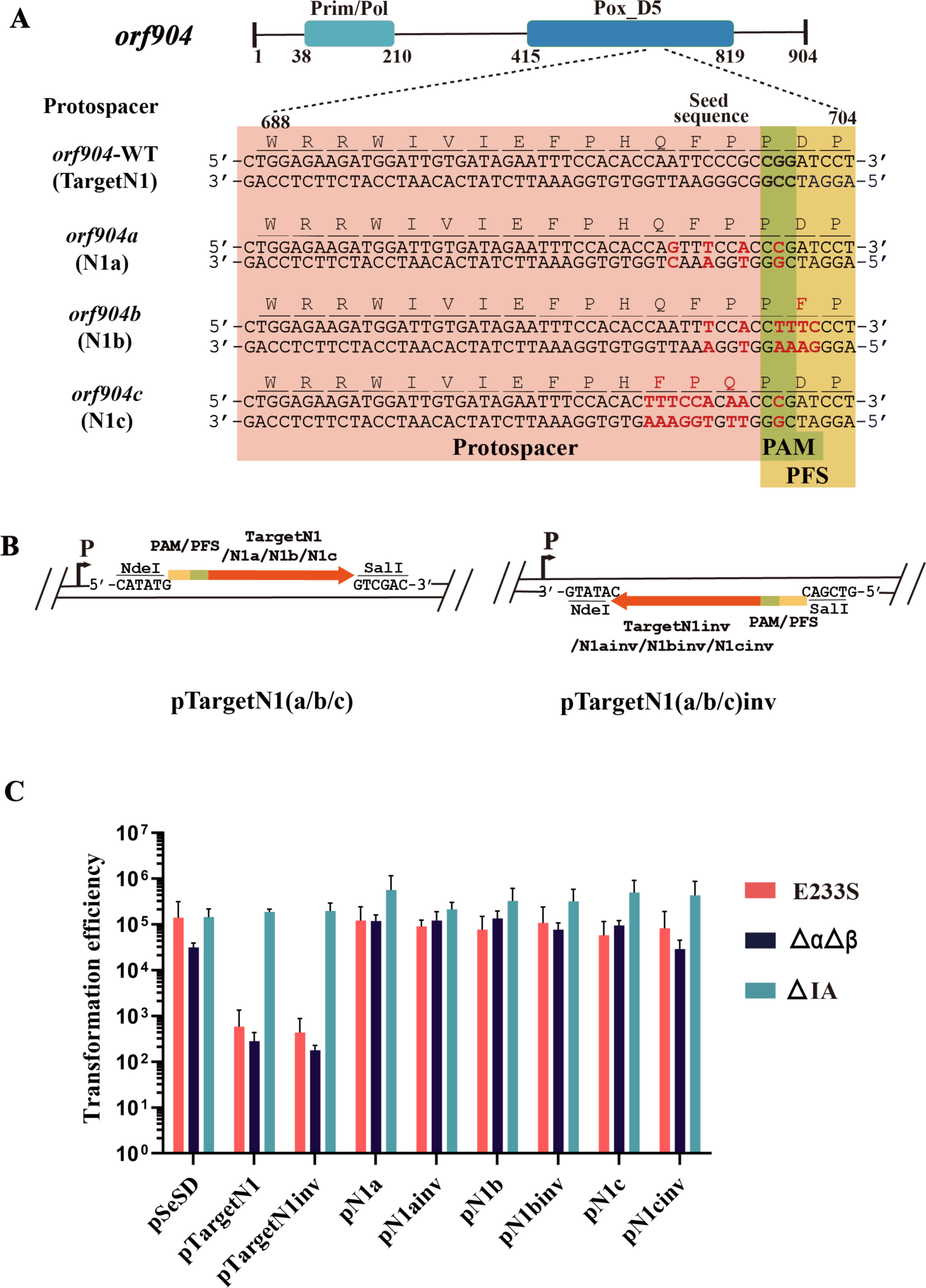
Rationale design of TargetN1 mutations and testing of their capability of evading the CRISPR immunization in *S. islandicus* REY15A. (A) The wild-type target sequence region and three mutated ones (N1a, N1b and N1c) are highlighted in the pink region whereas the flanking sequences are in the green and yellow region. Mutations both in DNA and in protein sequences are highlighted in red. (B) The design of protospacer modules in both orientation for the 3 derivative DNA segments based on the pRN1 N1a, N1b and N1c. (C) Transformation efficiencies of the corresponding protospacer/target plasmids. Three hosts were employed for transformation, including E233S: the wild-type host; ΔIA: E233S lacking the *cas* gene locus of IA CRISPR-Cas; ΔαΔβ: E233S lacking Cmr-α and Cmr-β loci.

These plasmids were then employed for testing their capability of inducing immune responses in the wild-type E233S strain and three mutants carrying deletions in distinctive sets of CRISPR elements (Table S1), including: (a) ΔαΔβ in which both III-B modules are removed, thus only carrying the I-A CRISPR system (37), (b) ΔαΔβΔarray, which is derived from ΔαΔβ by deletion of both CRISPR arrays (56); the immune system in this mutant should be inactive unless crRNAs can be provided from a CRISPR plasmid, and (c) ΔIA lacking *cas* genes of the I-A interference module (57). Transformation of the four archaeal hosts with pL2S56 yielded the following results: (a) while the reference plasmid pSeSD scored very similar rate of transformation (0.2-1×10^5^ colonies/μg), transformation of E233S and ΔαΔβ with pL2S56 or pL2S56inv triggered the IA immune responses, as the interference plasmid showed >100-fold reduced level of transformation (ca. 2×10^4^ colonies/μg), and (b) the immunity was abolished both in ΔIA and in ΔαΔβΔarray since each interference plasmid scored a transformation rate of ca. 10^6^ colonies/μg, which is comparable to that of the reference plasmid (Fig. 1C). The results showed that the plasmid-induced CRISPR immunity is dependent on the CRISPR arrays and the I-A CRISPR module but III-B CRISPR-independent. Very similar results were obtained with pTargetN1 and pTargetN1inv, indicating that the two mismatches between the pRN1 target and L2S56 do not influence the induction of the I-A immune responses. However, neither pL2S56inv nor pTargetN1inv could induce III-B responses in this archaeon (Fig. 1C). In a previous work, it has been shown that a relatively higher level of crRNA/target RNAs is required for triggering immunity of III-B CRISPR-Cas compared with the I-A systems (56), these data suggested that L2S56 crRNAs was expressed to a level that could be sufficient to elicit the I-A immunity but insufficient to trigger the III-B immunity for plasmid elimination in this archaeon.

### Identification of functional derivatives of pRN1 *orf904* that evade the L2S56-driven CRISPR immunity in the archaeal host

We then designed 3 derivative DNA segments based on the pRN1 target (Fig. 2A). The first mutant (N1a) possesses CGG at the PAM position and carries 3 additional mismatches at −2, −5 and −7 positions in the seed sequence region. The second mutant, denoted N1b, carries 6 nucleotide sequence changes including two mismatches at −2 and −5 positions in the seed sequence region and four changes at the PFS motif, yielding 5’-GAAAG-3’ motif at the target site, the last change makes a non-cognate target RNA upon transcription. The third one (N1c) has 9 mutations. Noticeably, while the designed mutations in N1a are synonymous, N1b and N1c have missense mutations: Those in N1b lead to the change of 1 amino acid and those in N1c, 3 substitutions (Fig. 2A). These DNA fragments were then cloned to pSeSD in both orientations (Fig. 2B), and the resulting plasmids were also tested for their ability of inducing CRISPR immunity in this archaeon using the interference plasmid assay. Specifically, three hosts were employed for transformation, including E233S, the wild-type strain; ΔαΔβ, the I-A CRISPR-active strain, and ΔI-A, the III-B CRISPR-active strain (Table S1). The results showed that, while TargetN1 induced the immune responses from the I-A CRISPR system, none of the 3 mutated targets, i.e., N1a, or N1b or N1c, were targeted by the CRISPR system in the archaeal host (Fig. 2C). Therefore, all designed protospacer mutations have successfully evaded the CRISPR immunity in *Sa. islandicus* REY15A, consistent with the results obtained with the interference plasmid assays shown in Fig. 1C.

Then, these mutated DNA segments were individually incorporated into *orf904* by inverse PCR with pN1SD as template and Gibson assembly of the resulting linear PCR fragments yielded pN1aSD, pN1bSD and pN1cSD, respectively. The synonymous mutations as well as the missense mutations carried by the plasmids are shown in Fig. 2A, Fig. S2. These plasmids were introduced into *S. islandicus* by electroporation to test the functionality of these pRN1 derivatives. As shown in Fig. 3, only a few colonies of transformants were obtained with pN1SD, indicating that the new wild-type pRN1-based shuttle vector was also restricted in *S. islandicus* RAY15A as seen for pREF11 in the same host (41). We also found that pN1cSD did not yield any colonies in transformation, suggesting that the carried missense mutations could have inactivated the replication protein, and this is also in agreement with the location of the target site in the conserved Pox_D5 domain of the pRN1 primase/polymerase (58, 59, 60). To this end, two mutated replicase genes (*orf904a* and *orf904b*) of pRN1 were found to be functional in the wild-type *Sa. islandicus* REY15A.

**Figure 3.**
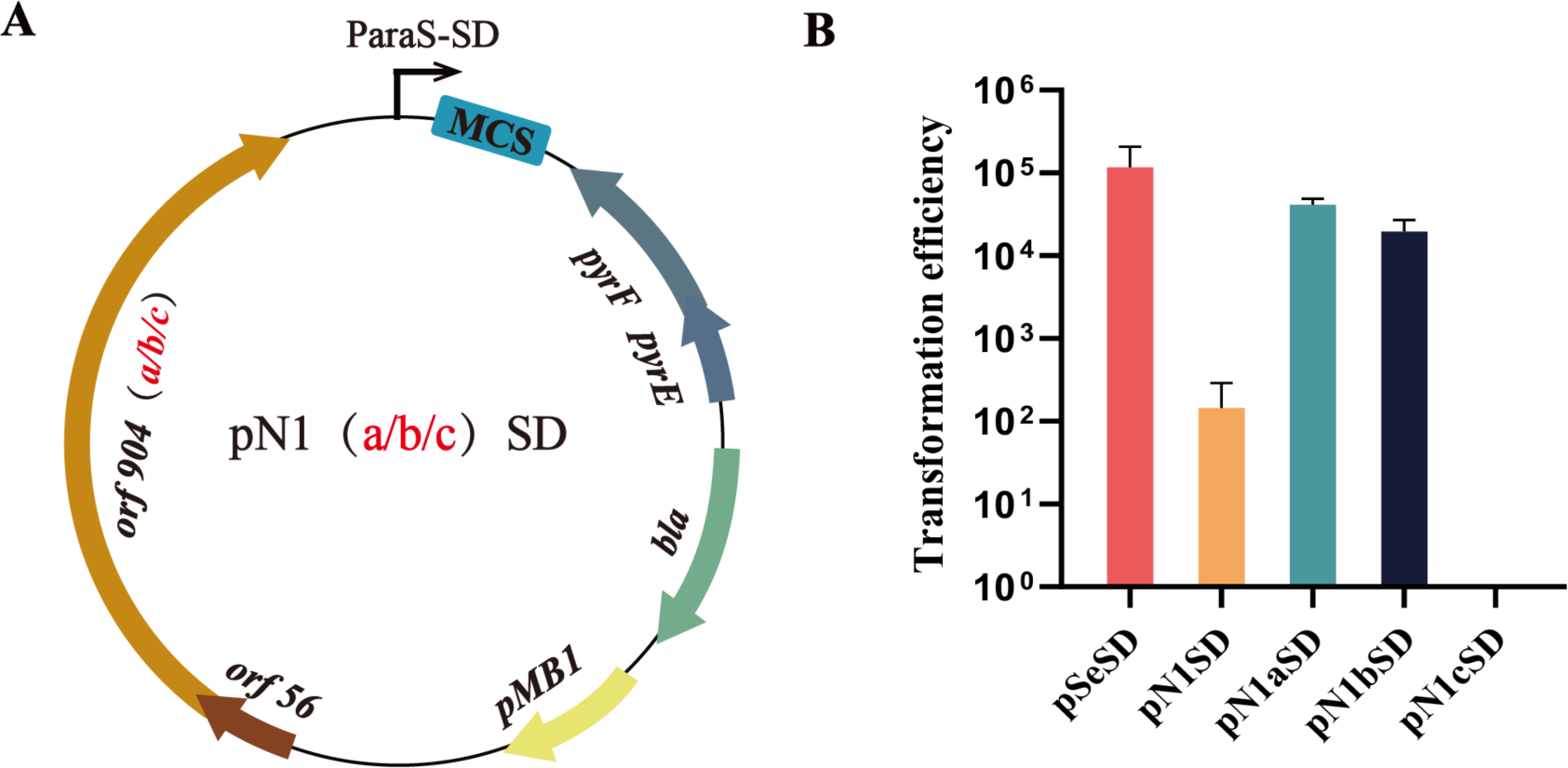
Functionality of the mutated *orf904* genes carrying a mutated pRN1 protospacer. (A) construction of vectors with each mutated *orf904* genes. (B) Transformation rates of the constructed pRN1-derived vectors.

### Testing the dual vector system with *Sa. islandicus pyrEF argD* double mutants

Previously, a *Sa. islandicus pyrEF argD* double mutant (E233SA, Table S1) was constructed and employed in testing the microhomology-mediated high-throughput gene inactivation in comparison with *Sa. islandicus* 16.4 (59). The genetic host can be selected with two genetic markers, one based on the complementation of *pyrEF* gene, while the other, on the *argD* complementation. Thus, it represents a suitable host for testing a dual plasmid system. A series of vectors were constructed in order to yield suitable vector for testing the dual plasmid system (Fig. S3). The first step was to construct a pRN1-derived shuttle vector with *argD* as the selection. This was done by amplifying an *argD* gene from *Saccarolobus solfataricus* with which the *pyrEF* marker in pN1aSD was replaced, producing pN1aBA. To facilitate the cloning with the vector, the NdeI in *orf904* was removed by reverse PCR and subsequent Gibson assembly, leading to pN1dBA. However, pSeSD and pN1dBA still share two elements, the pMB1 origin and the *bla* marker. To eliminate this redundancy, a different bacterial origin of replication and a kanamycin-resistant marker were amplified by PCR from p15AIE and pET30a, respectively. Assembling them with the *Saccharolobus* replicon of pN1dBA, yielded pN1dAA that only shares a promoter and a multiple cloning site with pSeSD (Fig. 4A-C, Fig. S3).

**Figure 4.**
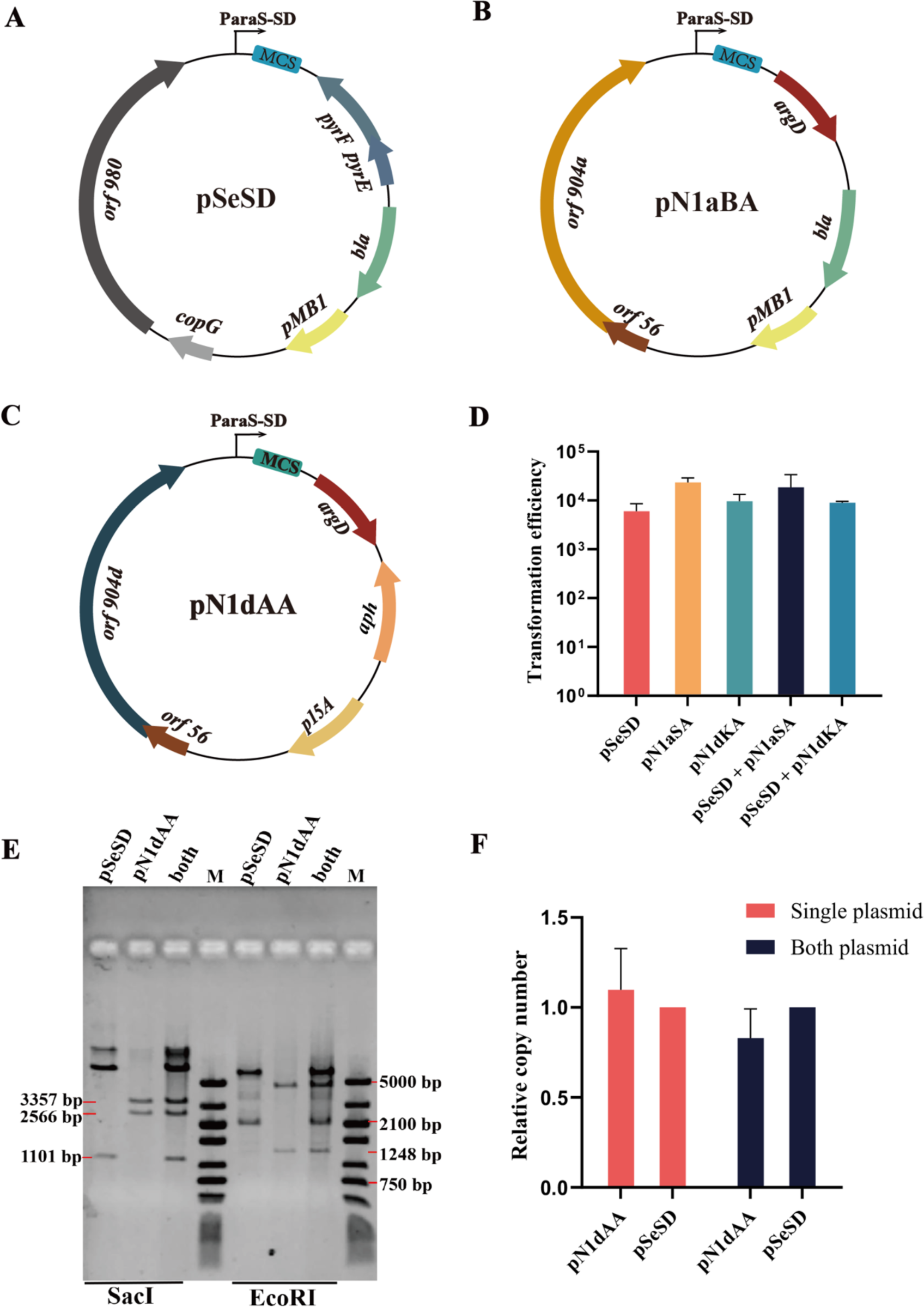
Compatibility of the pRN2- and pRN1-derived vectors in *S. islandicus*. (a-c). Maps of different plasmid vectors for *S. islandicus*. (D) Transformation rates with single plasmid or double plasmids. (E-F) Relative copy number of pSeSD (pRN2-derived) and pN1dAA (pRN1-derived) vectors. Gray values of DNA bands were quantified by the ImageJ software and normalized by the length of each fragment (kb). Relative copy numbers were then calculated with the copy number of pSeSD being set to 1.0 (F). Single plasmid: either pSeSD or pN1dAA; Both plasmids: pSeSD+pN1dAA; M: DNA marker ladder.

These vectors were then used to transform E233SA, either alone or in combination. As shown in Fig. 4D, transformation of *Sa. islandicus* cells either with each plasmid individually or with two plasmids simultaneously yielded very similar efficiencies. These results indicated these plasmid vectors are fully compatible with each other in the same cell, consistent with the concurrence of their wild-type plasmids in the original *Sa. islandicus* strain (21).

To test if the two plasmid vectors could be maintained at similar levels in host cells, strains carrying either pSeSD, or pN1dAA, or them both were grown in selective media to a late growth phase. Cell mass was then harvested for plasmid extraction. The resulting plasmid DNAs were then analyzed by restriction digestion and agarose gel electrophoresis. As shown in Fig. 4E, restriction fragments of predicted sizes appeared for each plasmid. Their relative ratio in the cell was determined by calculating the gray value of the enzymatic digestion products using the values of pSeSD as the reference (100%). This yielded the relative level of the two vectors in the single plasmid system was very similar to the cells carrying both plasmids (Fig. 4F). To assess plasmid maintenance in *Sa. islandicus*, these cultures were grown for 16 consecutive transfers (dilution of 20-25 folds was made in each transfer when OD600 values of these cultures reached 0.8-1.0). Plasmid DNAs were then prepared from the cell mass collected from the last batch of cultures and relative yields of plasmid were very similar to those prepared from the first batch of cultures as shown in Fig. 4E. These results showed both vectors are stably maintained in *Sa. islandicus* under the selection.

Sulfolobales organisms are widespread in hotsprings in the world (60). These organisms often coexist with extrachromosomal genetic elements, including viruses, cryptic and conjugative plasmids (61). In the CRISPR target database (62), reference sequences of plasmids and viruses available in the public GenBank databases are included, allowing CRISPR target sequences to be determined by searching the databases with sequences of CRISPR arrays. We conducted the search with the CRISPR arrays of *Sa. islandicu*s REY15A and found that several additional plasmids contain targets of the CRISPR spacers in this archaeon, including pANR4 conjugative plasmid exhibits the perfect match to Spacer 28 in Locus 1 (L1S28) of the CRISPR arrays. There are additional 3 conjugative plasmids, pING1, pKEF9 and pMGB1, each showing only 1 mismatch to L1S28, or L1S29 or L2S44 (Table S3), and these elements are also likely to be targeted in this archaeon. Indeed, investigation of pKEF9 conjugation in *Sa. islandicu*s revealed that the pKEF9-targeting spacer has been eliminated in conjugated cells via spacer deletion during conjugation (63). Moreover, 4 other plasmids (cryptic plasmid pHEN7, conjugative plasmids pARN3, pARN4, pAH1) exhibiting 5 mismatches to several spacers, e.g., L2S1 shows 5 mismatches to pARN3 and pARN4 (Table S3). Since III-B systems can tolerate mismatches between crRNAs and their targets (37, 53, 54, 55, 64), it would be interesting to investigate if some of these spacers could be targeted by the host CRISPR immune systems, and if so, whether they could be silenced differentially by the I-A and III-B CRISPR systems in this archaeon.

In conclusion, we have demonstrated that pRN1 is to be targeted by the IA CRISPR system of *S. islandicus* REY15A since it carries a DNA segment in *orf904* coding for the replicase of the plasmid, and design of pRN1 mutant escaping the host immunity by changing the CRISPR target sequence to a non-target one has yielded a fully functional plasmid backbone for vector construction.

## MATERIALS AND METHODS

### Strains and plasmids used in this work and cultivation

*Sa. islandicus* strains and plasmids used in this work are listed in Table S1. These archaea were grown in sucrose, casamino acids and vitamin (SCV) medium or in ACV in which sucrose is replaced with D-arabinose (19). If required, uracil and agmatine were supplemented to 2 μg/ml and 2 μg/ml, respectively, for cultivation of mutant strains. Incubation was at 75 °C with shaking at 200 rpm. Electroporation transformation of *Saccharolobus* cells was conducted with *ca.* 600 ng plasmid DNA, following the procedure previously described (19, 65).

### Enzymes and reagents employed in this work

All oligonucleotides employed in this work (Table S2) were synthesized in Tsingke (Qingdao, China), and restriction enzymes were purchased from ThermoFisher Scientific (Waltham, MA). Amplified PCR products were purified using a Cycle-Pure kit (Omega, Norcross, Georgia, USA). The designed plasmids were recovered by transformation of *E. coli*, with designed mutated sequences confirmed by DNA sequencing in Tsingke (Qingdao, China). Omega plasmid extraction kits for plasmid extraction were purchased from Omega, Norcross, Georgia, USA.

### Construction of target plasmids and plasmid interference assay

Five DNA fragments were designed carrying a wild-type or mutated target, i.e. the protospacer of the 56th spacer in locus 2 in the CRISPR array of the genetic host E233S (Proto-L2S56), its matching sequence in pRN1 (TargetN1) as well as 3 mutated targets (N1a, N1b and N1c) that contain mutations in the seed sequence region and/or mismatches in protospacer-adjacent motifs for the I-A CRISPR-Cas (45, 47) and protospacer-flanking sequences for the III-B immune systems (37, 53, 54, 55) in this crenarchaeon. Each DNA fragment was obtained by alignment of two overlapping oligos (Table S2) and subsequent polymerase extension. These DNA fragments were digested with LguI enzyme and cloned into pSeSD (34), yielding plasmid constructs pL2S56, pTargetN1, pN1a, pN1b and pN1c, which carries the wild-type L2S56 protospacer, target-N1 or each of the mutated targets (Table S1).

Interference plasmid assays were conducted as previously described (45, 53), and a great reduction in transformation rate with a test plasmid compared with the corresponding reference (>10 fold) is indicative of immune responses elicited by a target sequence on the plasmid (24).

### Construction of pRN1-based vectors

To construct pRN1-based *Saccharolobus-E. coli* shuttle vectors, a DNA fragment containing an *E. coli* replicon and the *pyrEF* selection marker (4196bp) was amplified from pSeSD (34) by PCR with primer pairs pSD-F/pSD-R, and the predicted minimal replicon of pRN1 (66), including the *orf56* and *orf904* genes, was amplified from pREF11 (41) by PCR using the primer pair of pREF-orf904-F/pREF-orf904-R. Subsequently, the two PCR fragments were circularized by the Gibson assembly (67) based on the *E. coli* RedET system (68) to yield pN1SD (Figure S1). The plasmid was then recovered by transformation of *E. coli* and confirmed by restriction analysis.

To generate pN1SD derivatives carrying mutated targets in *orf904* (see “Results and discussion”), primers N1a-F/N1a-R, N1b-F/N1b-R, N1c-F/N1c-R were designed and employed in inverse PCR reactions using Phanta Max Super-Fidelity DNA Polymerase (Vazyme Biotech, Nanjing, China) with pN1SD as the template. The resulting linear fragments containing N1a, N1b, or N1c DNA segment were self-circularized using the Gibson assembly to yield pN1aSD, pN1bSD and pN1cSD, respectively (Table S1, Figure S1). These plasmids were recovered by transformation of *E. coli* and their mutated targets were confirmed by DNA sequencing.

To construct pRN1 vectors with the *argD* selection, the marker gene was amplified by PCR from *Sa. solfataricus* P2 (69) using primer pairs P2-argD-F/P2-argD-R. In addition, a DNA fragment of pN1aSD lacking *pyrEF* was generated by inverse PCR using primer pairs pN1aSD-F/pN1aSD-R. The *argD* marker gene was fused together with the pN1aSD linearized fragment via Gibson assembly, giving the pN1aBA plasmid with the *bla* (beta-lactamase) marker in *E. coli* and the *argD* marker in *Sa. islandicus*. In addition, the occurrence of a NdeI restriction site in *orf904* prevents the utilization of NdeI in the multiple cloning sites region (MCS) for cloning. To remove that restriction site, primers N1-mNL-F and N1-mNL-R were designed and employed for NdeI site mutagenesis with SOE-PCR to yield pN1dBA (Figure S2).

To construct novel pRN1-based vectors, an *E. coli* replicon and selection marker different from those present on pSeSD should be used. To do that, DNA fragments containing the *E. coli* p15A origin were amplified by PCR from p15AIE using k15aori-F/15aori-N1-R primers whereas the *aph* gene encoding an aminoglycoside phosphotransferase was amplified from pET 30a using ak-F/k15aori-R primers. The third fragment carrying the *Saccharolobus* origin of replication and the *argD* marker was obtained by PCR from pN1dBA using 15aori-N1-F and ak-R primers. The resulting three fragments were recombined using the Gibson assembly reaction to yield the pN1dAA plasmid (Figure S1).

### Determination of the relative content of pRN1- and pRN2-derived plasmids

Relative plasmid content was determined by gel quantification of DNA bands after agarose gel electrophoresis. *S. islandicus* E233SA strains containing pSeSD, pN1dAA, and both of the above plasmids, respectively, were cultured to the OD_600_ of 0.8-1.0. Plasmid DNAs were extracted from cell mass using an Omega plasmid extraction kit. Equal volumes of plasmid preparations were digested with SacI or EcoRI respectively at 37°C for 1 h. The ImageJ software (70) was used to quantify the gray value ratio of digestion product bands. The resulting values were normalized by the length of DNA fragments, yielding relative gray values (pr. kb) that were used to calculate the relative copy number of pN1dAA and pSeSD plasmid in the cell.

## ACKNOWLEDGEMENTS

We thank members of the members of CRISPR and Archaea Biology Research Center at Shandong University for helpful discussions. This research was funded by National Key R & D Program of China, grant number 2020YFA0906800 to QS; National Natural Science Foundation of China (32270040 to QS, 32001022 to XF, 32370033 to YS).

## ETHANIC STATEMENT

This article does not contain any studies with human participants or animals performed by any of the authors.

## CONFLICT OF INTERESTS

The authors declare no conflict of interests.

## DATA AVAILABILITY

All relevant data have been reported in the submitted article, either in manuscript or in the supplementary data file.

## SUPPORTING INFORMATION

Additional Supporting Information for this article can be found online at xxxx.

## Supplementary figures and tables

**Figure S1.**
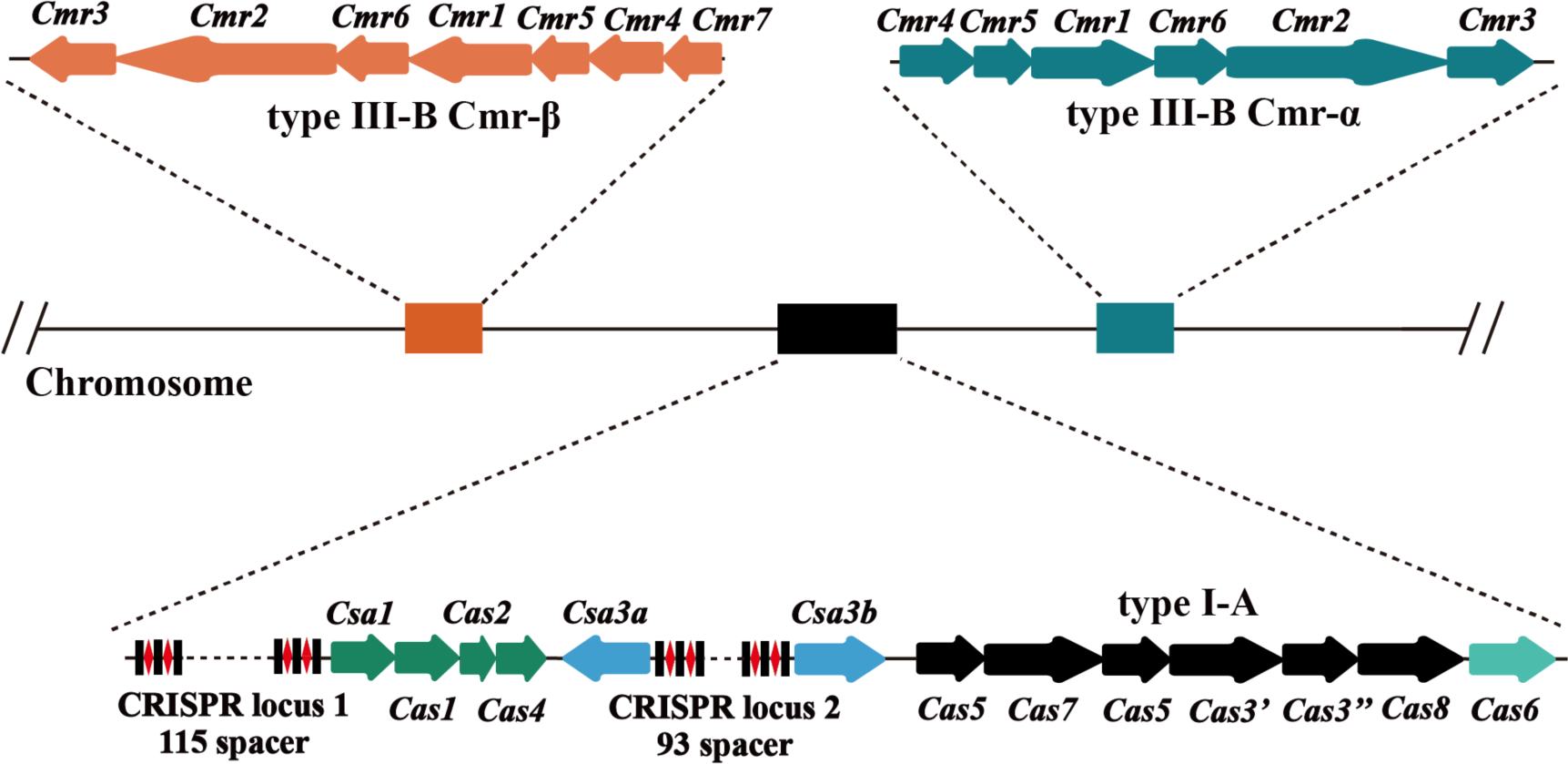
CRISPR-Cas systems encoded in *Sa islandicus* REY15A. Type I-A systems mediates the protospacer-adjacent motif-dependent DNA interference (1); Type III-B CRISPR systems (Cmr-α and Cmr-β) mediate the transcription-dependent interference of protospacer (2). The immunity is activated by cognate target RNAs showing mismatches to the 5’-repeat tag sequence of crRNA ( TGAAAG) (3, 4, 5), yielding invader clearance or cell dormancy or cell death (6, 7, 8). CRISPR locus 1 carries 115 spacers whereas locus 2 has 93 spacers. In addition, the host also carries a *cas* gene locus of adaptation (Csa1, Cas1, Cas2 and Cas4) responsible for spacer acquisition and two CRISPR-associated transcriptional regulators (Csa3a and Csa3b).

**Figure S2.**
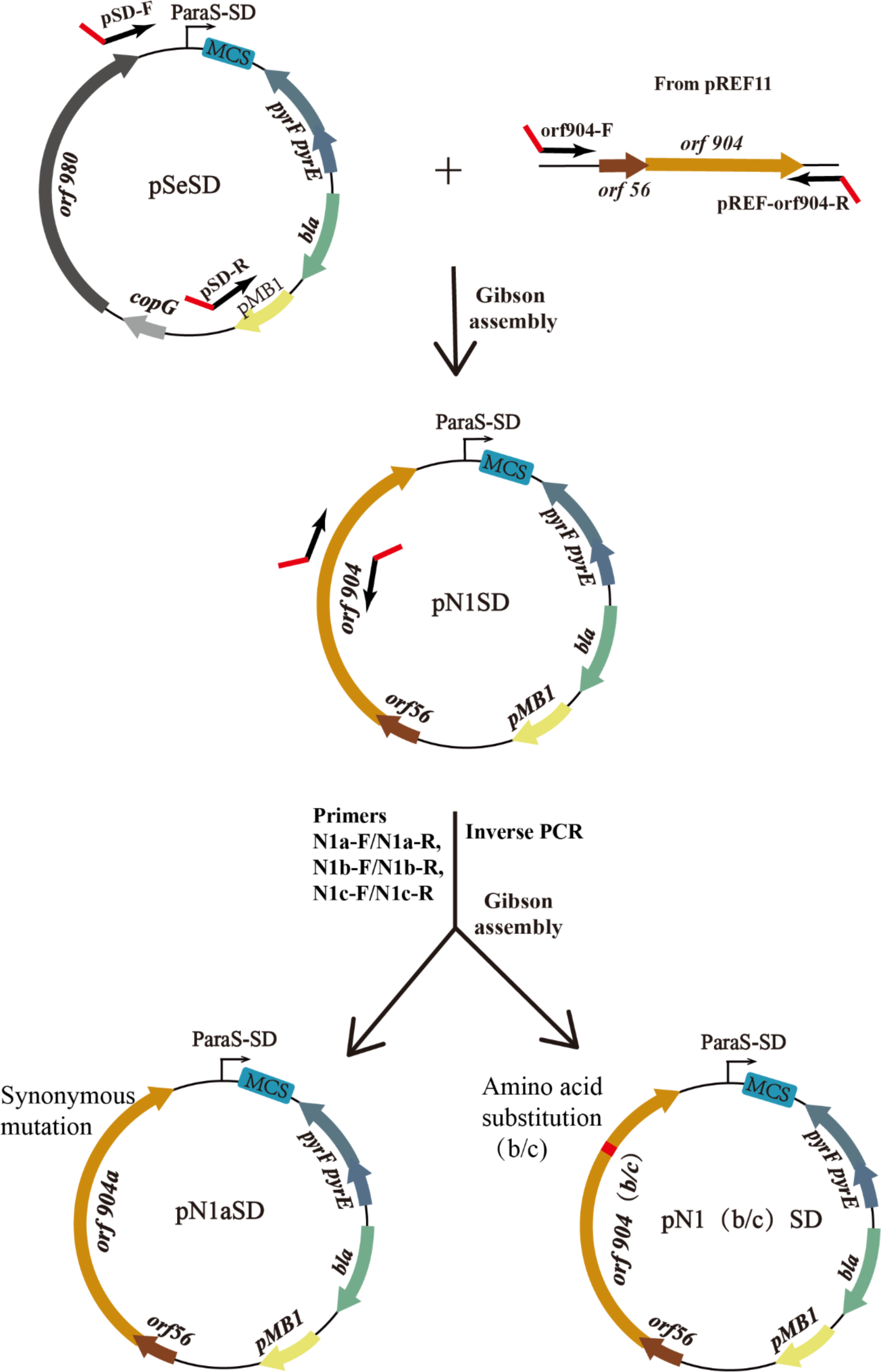
Construction of the pRN1-based *Saccharolobus-E. coli* shuttle vectors. A DNA fragment containing an *E. coli* replicon and the *pyrEF* selection marker (4196bp) from pSeSD plasmid and the predicted minimal replicon (including the *orf56* and *orf904* genes) of pRN1 from pREF11 were amplified with the primer pairs pSD-F/pSD-R and pREF-orf904-F/pREF-orf904-R, respectively. Subsequently, the two PCR fragments were circularized by Gibson assembly to yield pN1SD. To generate pN1SD derivatives carrying mutated protospacers in *orf904*, overlapping primers N1a-F/N1a-R, N1b-F/N1b-R and N1c-F/N1c-R containing the desired mutation were used for reverse PCR by using pN1SD as the template to introduce the *orf904a*, *orf904b* and *orf904c* into complementing plasmid pN1aSD, pN1bSD and pN1cSD, respectively. *pyrE*: Orotate phosphoribosyltransferase; *pyrF*: orotidine-5′-monophosphate decarboxylase; *bla*: beta-lactamase.

**Figure S3.**
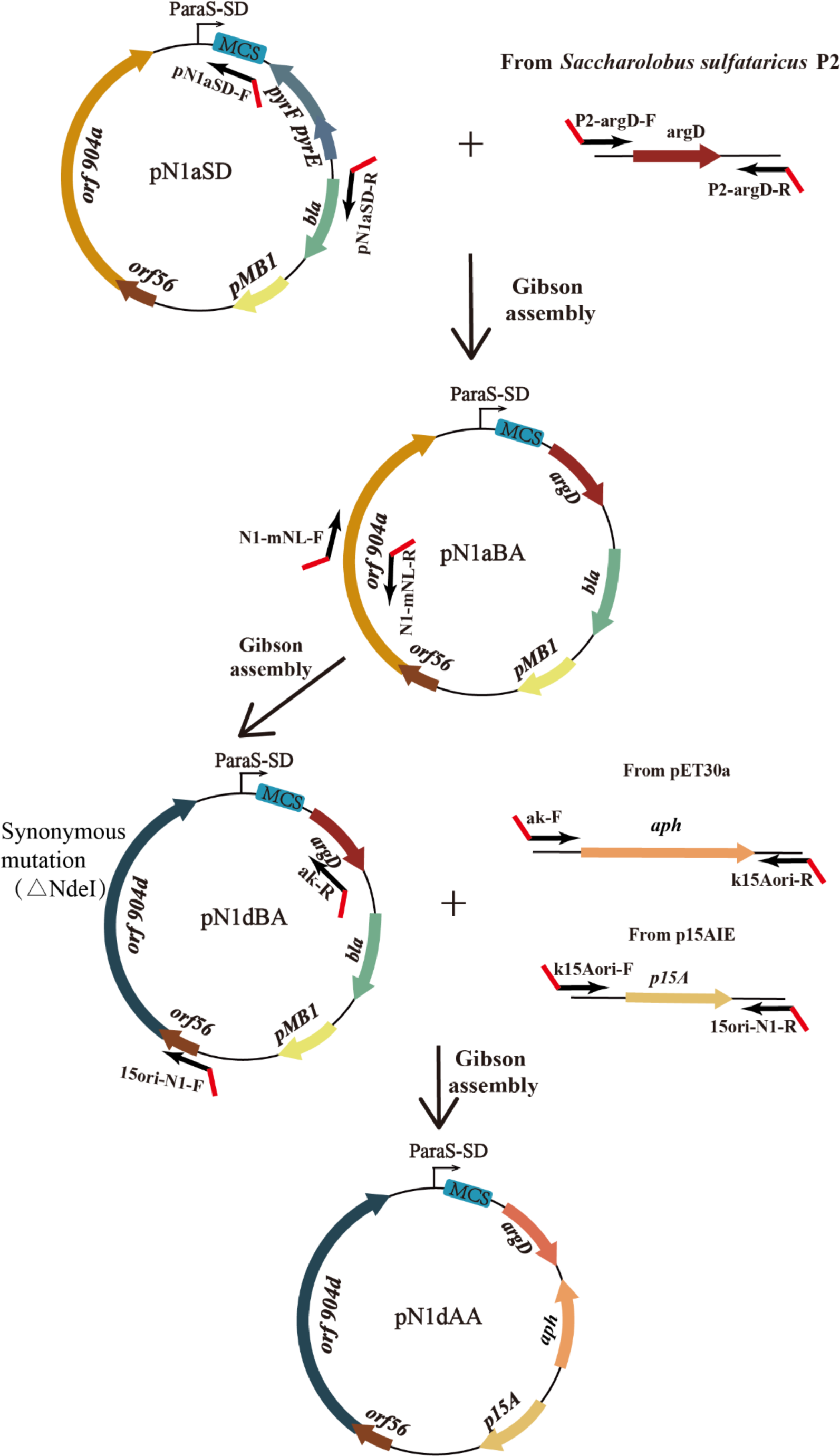
Construction of the pRN1-based *Saccharolobus-E. coli* shuttle vectors with an *E. coli* replicon and selection marker different from those on pSeSD. The *argD* selection marker gene from *Sa. solfataricus* P2 using primer pairs P2-argD-F/P2-argD-R was fused via Gibson assembly into the linearized pN1aSD lacking *pyrEF*, which was prepared by inverse PCR with the primer set pN1aSD-F/pN1aSD-R, to yield the pN1aBA plasmid. To remove the NdeI restriction site in *orf904*, primers N1-mNL-F and N1-mNL-R were designed and employed for NdeI site mutagenesis with SOE-PCR to yield pN1dBA. The *aph* selection marker gene amplified with k15aori-F/15aori-N1-R from pET30a and the p15aori amplified with ak-F/k15aori-R from p15AIE were combined with the PCR linearized pN1dBA plasmid using 15aori-N1-F/ak-R for Gibson assembly to form the final plasmid pN1dAA. *aph*: an aminoglycoside phosphotransferase gene; *argD*: an acetylornithine aminotransferase gene.

## Supplementary Tables

**Table S1.** *S. islandicus* strains and plasmids used in this work.

| Strain or plasmid* | Description | Reference |
| --- | --- | --- |
| *Strains-S. islandicus* |  |  |
| **E233S** | ***ΔpyrEF, ΔlacS*** | **(9)** |
| ΔIA-E233S | Derived from *S. islandicus* E233S, carrying deleting of *cas* genes of the I-A interference module | (10) |
| ΔαΔβ | Derived from *S. islandicus* E233S, carrying deleting of both III-B modules | (4) |
| ΔαΔβΔarray | Derived from *S. islandicus* ΔαΔβ by deletion of both CRISPR arrays | (11) |
| **E233SA** | ***ΔpyrEF, ΔlacS, ΔargD*** |  |
| *Plasmids* |  |  |
| **pSeSD** | ***A Sulfolobus-E. coli* shuttle vector derived from pRN2** | **(12)** |
| pREF11 | An unstable *Sulfolobus-E. coli* shuttle vector derived from pRN1 | (13) |
| pL2S56 | pSeSD carried a L2S56-protospacer | This work |
| pL2S56inv | pSeSD carried an inverted L2S56-protospacer | This work |
| pTargetN1 | pSeSD carried a pRN1-protospacer | This work |
| pTargetN1inv | pSeSD carried an inverted pRN1-protospacer | This work |
| pN1a/b/c | pSeSD carried the mutant pRN1-protospacer | This work |
| pN1a/b/c-inv | pSeSD carried h the inverted mutant pRN1-protospacer | This work |
| pN1SD | A pRN1-based *Sulfolobus-E. coli* shuttle plasmid, consisting of an orf56-orf904 fragment of pRN1 fused to a 0-4196 bp fragment of pSeSD, carrying the *pyrEF* gene | This work |
| pN1a/b/cSD | Mutation of the *orf904* sequence in the pN1SD | This work |
| pN1aSA | Replace *pyrEF* gene with *argD* gene in pN1aSD | This work |
| pN1dSA | Mutation of the NdeI site of *orf904* in pN1aBA | This work |
| **pN1dKA** | **Replace *Amp<sup>r</sup>* gene with *Kan<sup>r</sup>* gene, and *pMB1* with *p15A-ori* in pN1dBA** | **This work** |

\*Genetic hosts and plasmid vectors for *Sa islandicus* REY15A are highlighted in bold face.

**Table S2.**
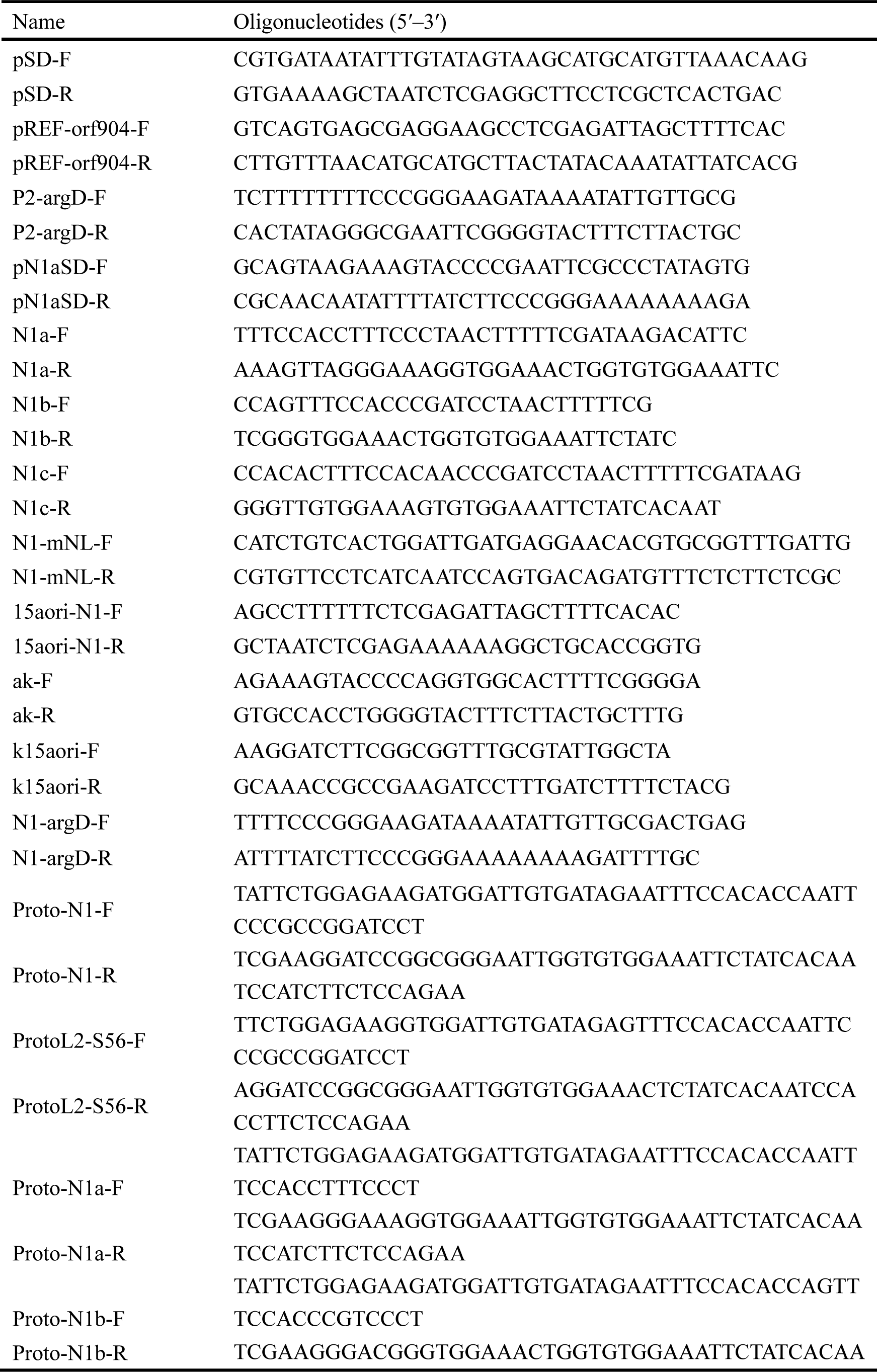
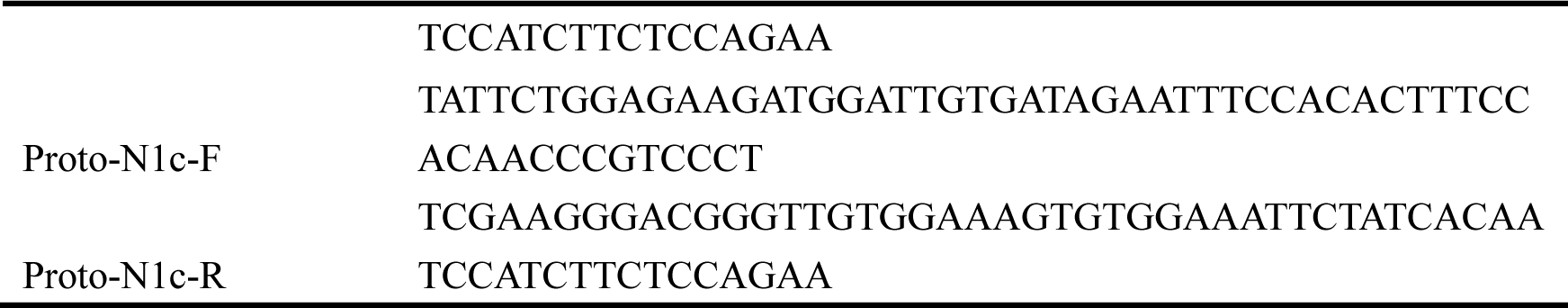
Oligonucleotides and primers used in this work.

**Table S3.**
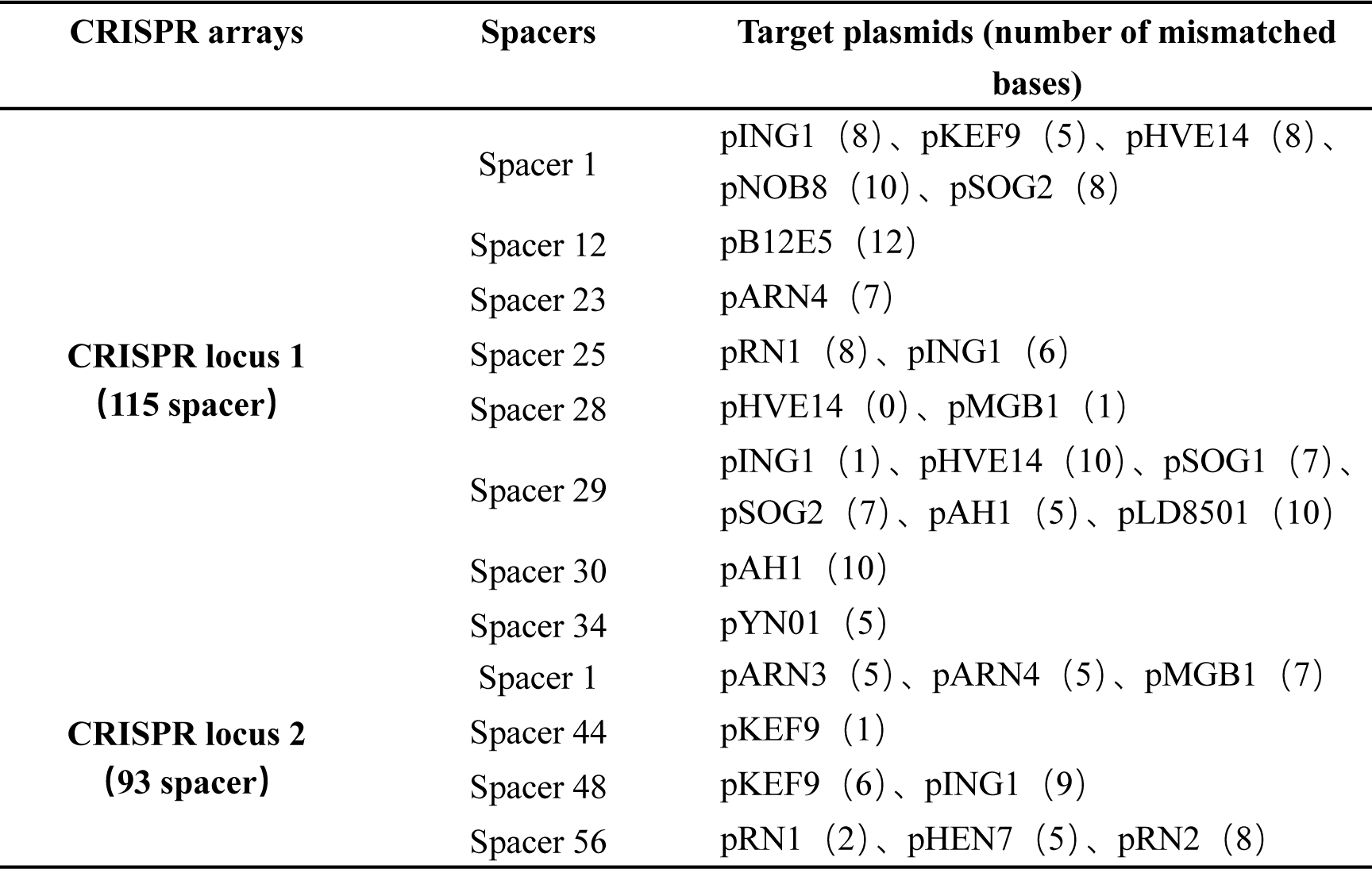
Plasmid-matching spacers in the genome of *S. islandicus* REY15A.

## Notes

### Competing Interest Statement

The authors have declared no competing interest.

